# 3D Variability Analysis: Resolving continuous flexibility and discrete heterogeneity from single particle cryo-EM

**DOI:** 10.1101/2020.04.08.032466

**Authors:** Ali Punjani, David J. Fleet

## Abstract

Single particle cryo-EM excels in determining static structures of protein molecules, but existing 3D reconstruction methods have been ineffective in modelling flexible proteins. We introduce 3D variability analysis (3DVA), an algorithm that fits a linear subspace model of conformational change to cryo-EM data at high resolution. 3DVA enables the resolution and visualization of detailed molecular motions of both large and small proteins, revealing new biological insight from single particle cryo-EM data. Experimental results demonstrate the ability of 3DVA to resolve multiple flexible motions of *α*-helices in the sub-50 kDa transmembrane domain of a GPCR complex, bending modes of a sodium ion channel, five types of symmetric and symmetry-breaking flexibility in a proteasome, large motions in a spliceosome complex, and discrete conformational states of a ribosome assembly. 3DVA is implemented in the *cryoSPARC* software package.

## 1 Introduction

Many protein molecules have a mechanistic function, exhibiting flexible motion across a continuous landscape of energetically favourable conformations. Single particle cryo-EM collects thousands of static 2D protein particle images that, in aggregate, span the target protein’s 3D conformational space. As such, cryo-EM holds great promise for studying proteins with variation in conformation, and in turn, protein dynamics and function [23]. However, there are significant computational challenges in resolving continuous conformational variability from static image data, stemming from the need to simultaneously estimate the canonical structure of the molecule, the multiple ways in which the structure can change (due to motion, dissociation, etc), and the state of the molecule in each particle image. These problems are exacerbated by high levels of image noise.

Given a sample for a heterogeneous target molecule, the 3D structures corresponding to the particle images lie on a manifold in the abstract space of all possible 3D structures. The manifold may have a non-linear geometry in general, but for many molecules of interest a linear subspace model provides an effective way to represent the 3D structures in a cryo-EM sample. A linear subspace model represents 3D structures in terms of a 3D base structure plus a weighted sum of 3D basis functions. The weights represent the conformational state of the molecule in a given particle image. Subspace models, such as Principal Component Analysis (PCA), have been discussed extensively in the cryo-EM literature [1, 31, 46, 48]. They are attractive for their simplicity, analytical properties, and the existence of stable numerical methods for eigenvector (basis function) computation, but they are not widely used in practice (e.g., compared to discrete 3D classification [13, 40, 42, 43]). The primary barriers have been the lack of efficient methods for estimating high resolution 3D models from 2D image data, especially when the observed images are projections from unknown orientations, and modified by different contrast transfer functions.

We introduce *3D Variability Analysis* (3DVA), a method for fitting 3D linear subspace models to single particle cryo-EM data. Following Tagare et al [46], the algorithm is formulated as a variant of the Expectation-Maximization algorithm [8, 29] for Probabilistic PCA [39, 47]. Thee use of Expectation-Maximization facilitates the estimation of 3D models from 2D data, and by careful formulation of the numerical optimization, we obtain a efficient method for computing high-resolution representations. Through experimental results we show that 3DVA allows for directly resolving molecular motion, flexibility, changes in occupancy, and even discrete heterogeneity of a wide range of protein molecules. It resolves high-resolution flexibility, with motions of individual *α*-helices as small as a few angstroms, for both large and small proteins. To our knowledge, this is the first method to resolve flexible conformational variability of small proteins, such as the distinct types of flexible motion we find in the sub-50 kDa transmembrane region of a GPCR complex, at resolutions below 4Å.

By resolving continuous conformational changes, 3DVA reveals biological insights from single particle cryo-EM data beyond the reach of existing methods, simplifying the analysis of conformational heterogeneity. We show that it can resolve ratcheting motions of a ribosome, large flexible motion of a pre-catalytic spliceosome complex, and several discrete conformational states of a bacterial large ribosomal subunit assembly. For such large, multi-MDa complexes, recently proposed deep generative models [53, 52] have been experimentally demonstrated to resolve similar continuous and discrete heterogeneity at coarse resolution. However, 3DVA is 20-50x faster to optimize, requiring no manual parameter tuning or hyperparameter optimization on individual datasets.

A GPU implementation of 3DVA in *cryoSPARC* allows one to process large experimental datasets with 10^6^ particle images. It resolves multiple variability components at resolutions as high as 3Å-4Å, limited only by GPU memory and signal present in the data. 3DVA was released in the *cryoSPARC* [34] software package v2.9+ and has been used in several structural studies, including the SARS-CoV-2 virus spike protein [50], mTORC1 docked on the lysosome [38], human oligosaccharyltransferase complexes OST-A and OST-B [36], bacterial unfoldase-protease complex ClpXP [37], a viral chaperonin [45], the *Acinetobacter baumannii* 70S Ribosome [26], and Adrenomedullin Receptors AM1 and AM2 [22].

## 2 Linear Subspaces, PCA, and 3D Variability Analysis

Under the standard cryo-EM image formation model [9, 14, 33, 41], a 2D particle image, *X_i_*, is a corrupted projection of the target 3D density 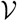 from an unobserved pose *ϕ_i_*, plus additive noise, *η*. As is common in cryo-EM formulations, we represent 3D densities, 2D images, the CTF and projection operator in the Fourier domain. For example, for a box-size with linear dimension *N*, 3D density maps are represented as 3D grids of *N*^3^ Fourier coefficients. Accordingly, we can formulate the standard generative model as follows:

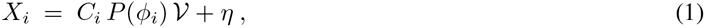

where *P* (*ϕ_i_*) is the projection operator and *C_i_* is the contrast transfer function (CTF) for image *i*. This model assumes all images are generated from a single 3D density map 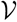. It assumes a homogeneous population of molecules, all in the same conformation, differing only by a rigid coordinate transformation. Discrete 3D classification methods [13, 40, 42, 43] extend this model, assuming each particle image is generated from one of a small number of of distinct, independent 3D densities. Nevertheless, methods based on homogeneity or discrete heterogeneity are of limited value for samples with continuous heterogeneity, as they yield low quality reconstructions.

3D variability analysis accounts for continuous conformational flexibility interms o f a *K*-dimensional linear subspace model. Under a variant of probabilistic PCA [39, 47], each experimental particle image is generated from a **mean density**, *V*_0_, plus weighted contributions of *K* **variability components**, *V*_1_ through *V_K_*, denoted by *V*_1:*K*_. The model also includes a per-particle scale factor, *α_i_*, to account for varying ice-thickness and contrast level and additive Gaussian noise, *η*. The noise is assumed to be zero-mean with a diagonal covariance matrix in the Fourier domain, with constant variance in annular rings. Formally, in the Fourier domain, the generative model becomes

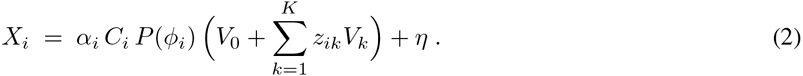

Each *V_k_* is a 3D density represented in the Fourier domain, capturing a particular type of change to *V*_0_. The vector of per-particle weights, **z**_*i*_ = (*z_i_*_1_*, …, z_iK_*) are also known as the **latent coordinates**.

The predominant method for fitting linear subspace models in many domains is the Principal Component Analysis (PCA) algorithm [3], under the assumption that the basis functions are orthogonal; i.e., 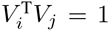 if *i* = *j*, and 0 otherwise. Given a random data sample (e.g., 3D densities), assumed to be generated independently from the same underlying distribution, for PCA one would compute the sample mean (*V*_0_ in Eq. (2)), and the leading eigenvectors of the covariance matrix of the data distribution (i.e., those with the largest eigenvalues). These *principal directions* account for the most significant factors of variation in the data (like *V*_1:*K*_ in Eq. (2)).

Direct application of PCA to single particle cryo-EM data is challenging however. First, each 2D image is a partial observation of the 3D density from one direction, so construction of the full covariance of 3D densities from images is non-trivial. Second, the high dimensionality of the covariance matrix means that it may not be possible to store in memory, or to compute its eigenvectors except at low resolutions. Third, each image is modified by a different CTF, violating the PCA assumption that data are generated from the same distribution.

To mitigate these challenges one could restrict analysis to 2D by computing the 2D covariance of particles within a single 2D class or viewing direction (cf. EMAN2 [2]). This removes the partial observation problem, but fails to capture 3D variability or account for the CTF [48]. In 3D one can use bootstrapping [31], wherein multiple random subsets of images are used to reconstruct multiple 3D structures under the homogeneous model (Eq. 1). With such “bootstrapped” 3D densities one avoids the missing information problem due to projection, but it can only operate at low resolutions to avoid high-dimensional covariance matrices. It further suffers from statistical inefficiency due to the random noise injected into each bootstrap sample. Together, these drawbacks limit such methods to resolving coarse resolution eigenvectors that capture gross structural changes of large protein complexes [31]. One can also employ low-dimensional approximations to the covariance, which mitigate storage and computational cost, but such methods have been demonstrated only at coarse resolutions [1].

As explained in Methods, 3D Variability Analysis overcomes these issues with conventional PCA techniquees, enabling subspace models at high resolution and for smaller proteins. Inspired by Roweis [39], we formulate 3DVA as a form of Probabilistic PCA, assuming that data are drawn from a high dimensional Gaussian distribution, with Gaussian observation noise in Eq. (2), and a Gaussian prior over latent coordinates. Following [39], the Expectation-Maximization algorithm [8, 29] is used to obtain a Maximum Likelihood (ML) estimate of the variability components *V*_1:*K*_. Crucially, this PPCA algorithm accommodates partial observations (e.g., 2D projections of 3D structures), and data-specific corruption (e.g., the CTF in cryo-EM data), and it works well with high-dimensional data without the need to explicitly store or approximate the covariance matrix [39]. It also supports direct estimation of only the top *K* components, as desired in 3DVA.

The 3DVA algorithm assumes the mean density, *V*_0_, along with the per-particle CTFs, *C_i_*, and poses, *ϕ_i_*, are known. These quantities could be estimated by processing the image data using standard homogeneous refinement. Given these quantities, and random initial values for the variability components, 3DVA uses iterative optimization, each iteration of which has an E-step and an M-step. Following [39, 29], the E-step updates the mean of the posterior distribution over the latent coordinates, **z**_*i*_, independently for each image *X_i_*. The M-step then uses the expected log likelihood to update the per-particle scale parameter *α_i_*, and the variability components, *V*_1:*K*_.

One issue with such optimization problems is the computational cost associated with estimating large numbers of unknowns, given large numbers of image observations. As explained in Methods, we show how to take advantage of the problem structure, decoupling the estimation problem into smaller subproblems that can be solved much more efficiently. In addition, during optimization, the variability components *V*_1:*K*_ are regularized by a low-pass filter and high-pass filter. The low-pass filter attenuates high-frequency noise to prevent over-fitting. An optional high-pass filter removes sources of low-resolution variability caused by contaminants (e.g. denatured protein mass at air-water interfaces). Beyond these optional parameters, and the specification of the resolution limit and the number of variability components, 3DVA does not require tuning or parameter changes between datasets.

When 3DVA is applied to single particle EM data, it outputs variability components, *V*_1:*K*_, and per-particle latent coordinates, **z**. Variability components are those directions in the space of 3D density that, from the mean structure, *V*_0_, define a linear subspace that best fits the principal directions of variation in the data. Each component can be understood as a 3D voxel grid of density values (computationally represented in Fourier space). These voxel values indicate where density should be added or removed to explain variability amongst the particles. Orthogonality of the *K* components ensures that each explains a different mode of variation. Fig. 1 shows an example of a variability component on a ribosome dataset. Variability components found by 3DVA often capture flexible motion of the underlying molecule in a sample. Fig. 1d,e show qualitatively how linear subspace models capture object motion. Displacements up to the scale of the object are well approximated. Small objects moving large distances are less well modelled. That is, 3DVA can represent large motion in low resolution regions of a 3D density map, as well as small motions in high resolution regions.

**Figure 1:**
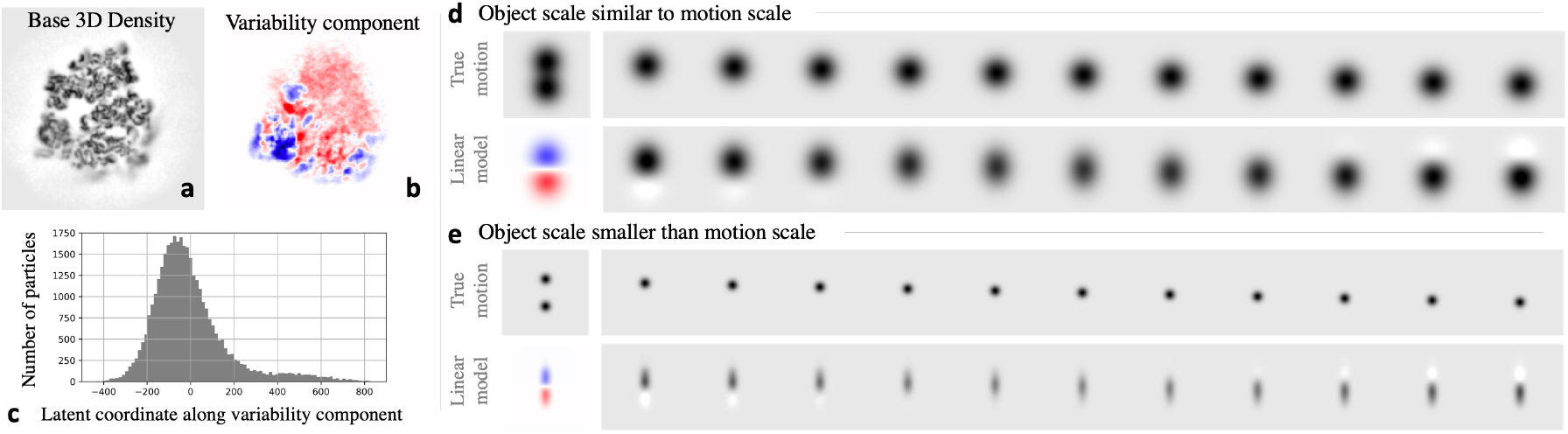
Examples of 3DVA. **a:** A central slice through a 3D density map reconstructed from 80S ribosome particle images. **b:** A central slice of the first 3D variability component (transformed to real-space) for the reconstruction in **a**. Positive (red) and negative (blue) values correspond to density to be added and subtracted from the base density in **a** to explain heterogeneity in the data. This variability component captures motion of the small subunit (bottom left, darker blue) relative to the large subunit. **c:** The distribution of latent coordinates for the component in **b**. **d, e:** Synthetic examples of how a linear subspace model represents object motion. In both cases, the top row shows observed images of an object in motion, while the bottom shows the first variability component (in red / blue) and images generated by the model to explain the motion. When the displacement is small relative to object size (**d**) the model approximates the motion well. When the object is smaller than the motion (**e**), the linear model is less accurate.

3DVA also yields latent coordinates, **z**, for each particle image. They indicate the level to which each variability component is present in a given image. If the *i*th particle image has a large positive value for *z_ik_*, this means it is best described by adding a large amount of *V_k_* to *V*_0_. The variance of each latent coordinate across the images provides a measure of importance, since components with large variance explain the most variability in the data. When a population of particle images contains well-separated clusters of different conformations, 3DVA components identify differences between clusters, and the latent coordinates of particles will be similarly clustered, providing insight into discrete heterogeneity.

## 3 Results

In what follows, we consider applications of 3DVA to one synthetic dataset and six real-world cryo-EM datasets. Results on synthetic data help to validate the approach and convergence of 3DVA to the true underlying principle linear subspace of the data. Results on the real-world datasets demonstrate the ability of the method to resolve high resolution continuous and discrete conformational changes from single particle cryo-EM.

For the real-world experimental data we first compute a consensus refinement using all particle images. In the case of membrane proteins, non-uniform refinement [35] is used to improve image alignments and overall resolution. The resulting particle images and pose alignments are then used to compute variability components and latent coordinates via 3DVA. No prior information is provided about the type or form of heterogeneity in each dataset. 3DVA is run with a real-space mask (see Methods) that excludes solvent, and for membrane proteins the mask also excludes detergent micelle or lipid nanodisc. With all experimental datasets here, 3DVA executes 20 iterations of Expectation-Maximization, starting from random initialization. All experiments are run on a single workstation with one NVIDIA V100 GPU.

For data with continuous heterogeneity, we render the 3D density generated by the subspace model at positive (blue) and negative (red) values of the latent coordinate for individual variability components, along with a superimposed rendering of both densities to clearly indicate the difference between them. Guide lines (solid and dashed) and feature markers (+) provide visual reference points, allowing visual comparison of the 3D maps from positive and negative latent values. For each dataset, videos displaying flexible conformational changes are available as supplementary movies.

### 3.1 Synthetic T20S

We generate synthetic data by deforming the 3D density map of a T20S proteasome [4] at 3*Å*. The *ground truth* deformations lie in a three dimensional linear subspace, the basis functions for which correspond to bending along each of the *x*, *y*, and *z* axes (Figure 2).

**Figure 2:**
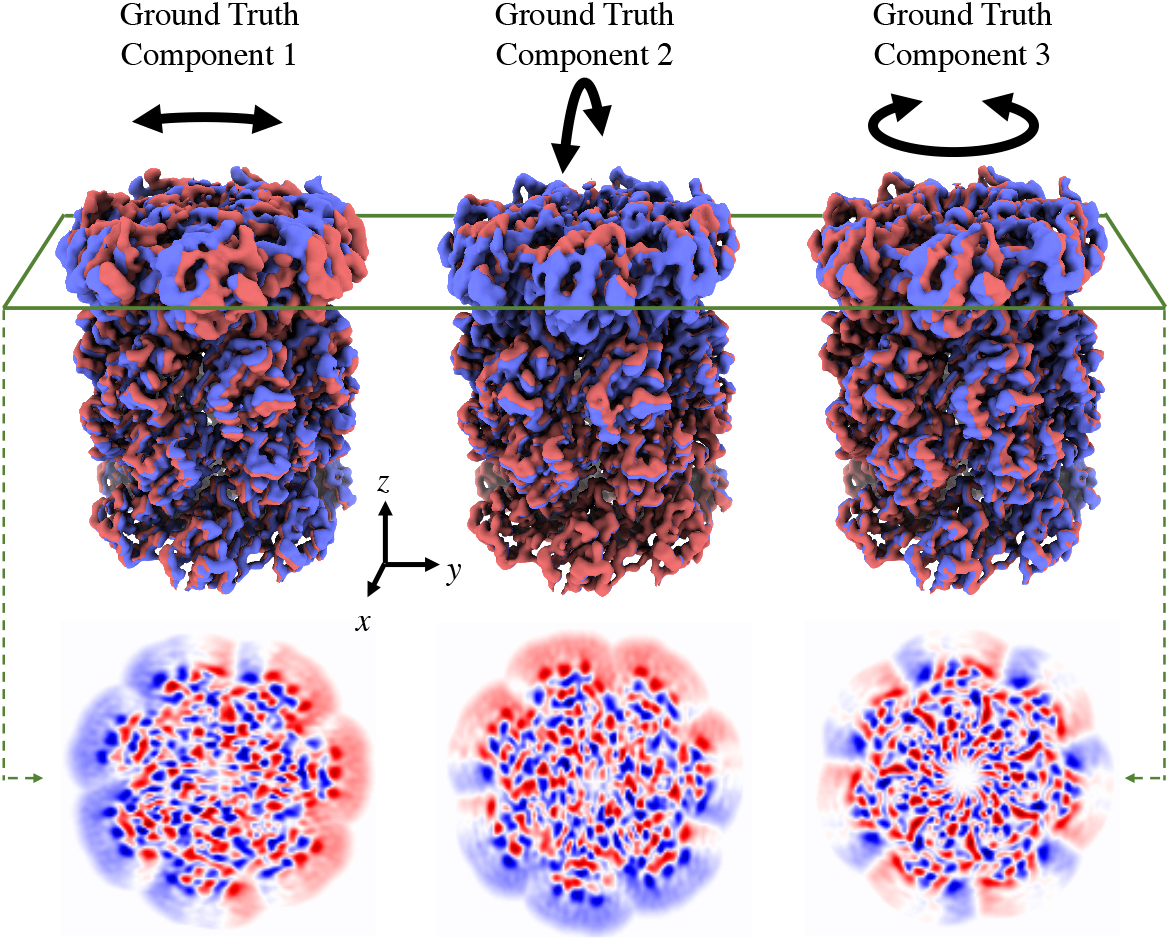
Ground-truth linear subspace directions of a synthetic T20S proteasome molecule used to generate synthetic particle image data. **Top row:** 3D renderings of the consensus density plus (red) and minus (blue) each subspace direction at one standard deviation. The three components represent bending of the molecule around the *x*-axis, bending around the *y*-axis, and twisting about the *z*-axis. **Bottom row:** Slices in the *x – y* plane of the three subspace directions depicted in the top row. The slices show positive and negative values where density is added and subtracted according to the subspace direction.

Each synthetic dataset comprises 100,000 particle images. For each particle image, we randomly sample *ground-truth* latent coordinates from a mean-zero Gaussian distribution with a diagonal covariance matrix, the diagonal variances of which are 50^2^,40^2^ and 30^2^, corresponding to bending along the *x*, *y*, and *z* axes. The corresponding deformation is then applied to the T20S density map. A viewing direction is then randomly drawn from a uniform distribution over the sphere. For CTF parameters, average defocus is drawn at random uniformly from the interval [0.5*μm,* 2.0*μm*], astigmatism from the interval [−0.1*μm,* 0.1*μm*], and astigmatism angle uniformly around the circle. Accelerating voltage is set to 300kV, spherical aberration to 2.7mm, and per-particle scales to 1.0. Particle images are generated using Eq. 2.

Particle images are collectively normalized to have unit signal variance on average. White noise is them generated and added to the particle images. To this end we use four noise levels, namely, *σ* = 0, 4, 10, and 20, corresponding to noiseless data, and SNRs of 1/16, 1/100, and 1/400 respectively. This yields four datasets, each with 100,000 particle images, that are used to test 3DVA (see Fig. 3a).

**Figure 3:**
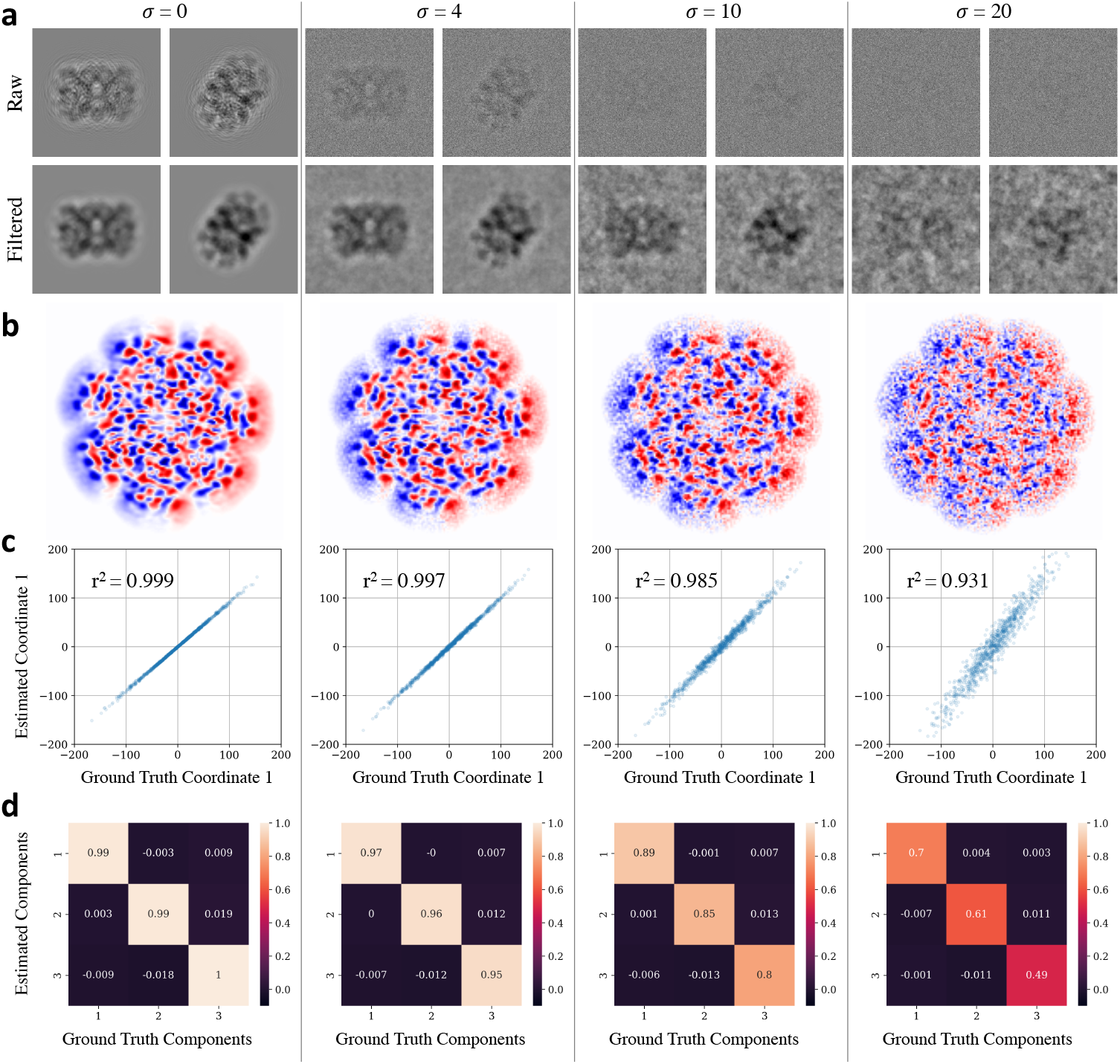
Synthetic data and results of 3DVA with four noise levels, i.e., *σ* = 0, 4, 10, and 20. 3DVA is run with particles at each noise level, solving for three variability components. **a:** Example particle images are shown, with raw images on top with low-pass filtered versions below (cut-off at 30*Å*). In the noiseless case, ringing from the CTF is visible in the raw images. As the noise level increases, the particle become harder to distinguish, even after filtering. **b:** Results of 3DVA used to recover variability components. Slices in the *x − y* plane through the first variability component are shown, with red and blue corresponding to positive and negative values. These slices can be compared with ground truth slices in Fig. 2 (bottom left). **c:** Scatter plots showing ground-truth latent coordinates for component 1 (*x*-axis) vs 3DVA estimated latent coordinates for component 1 (*y*-axis) for 1% of the images in the dataset, along with correlation coefficients (*r*^2^ values). **d:** Confusion matrices indicate alignment between ground-truth and 3DVA variability components in terms of the correlation coefficient between true (*x*-axis) and estimated (*y*-axis) components. 3DVA resolves the true subspace directions in the correct order from most to least significant, with nearly zero confusion between the subspace directions. Correlations decrease as increasing noise.

At each noise level, we run 3DVA using the ground-truth poses and CTF parameters for each particle, estimating three variability components. A consensus reconstruction of input particles is used as the mean density *V*_0_. Low-pass filtering is not used, and initialization is random. In all cases, 3DVA correctly recovers the true subspace (Figure 3b), as well as the latent coordinates of particles (Figure 3c). Noise is present in the estimated variability components due to the inherent noise in the input images. When the noise level is low or zero, latent coordinates are recovered with nearly perfect correlation to the true coordinates. As noise increases the estimates become more varied, but correlation with the true coordinates remains high (Figure 3c). In all cases, 3DVA recovers the three subspace directions in the correct order, and without significant confusion between the directions (Figure 3d). In the noiseless case, correlations are nearly perfect. In the three noisy cases, correlation decreases as the variability components are affected by noise.

Despite the noise in input images, 3DVA estimated variability components do recover the true signal in the sub-space directions at high resolutions. To see that this is the case, we first compute FSC curves between the consensus reconstruction using particles at each noise level and the ground-truth consensus reconstruction (Figure 4a). This baseline indicates the degree to which noise limits the algorithm’s ability to recover signal from the finite number of images. As expected, the noiseless reconstruction has nearly perfect correlation at all frequencies against the ground-truth. As the noise level is increased, resolvable signal decreases and resolution is limited.

**Figure 4:**
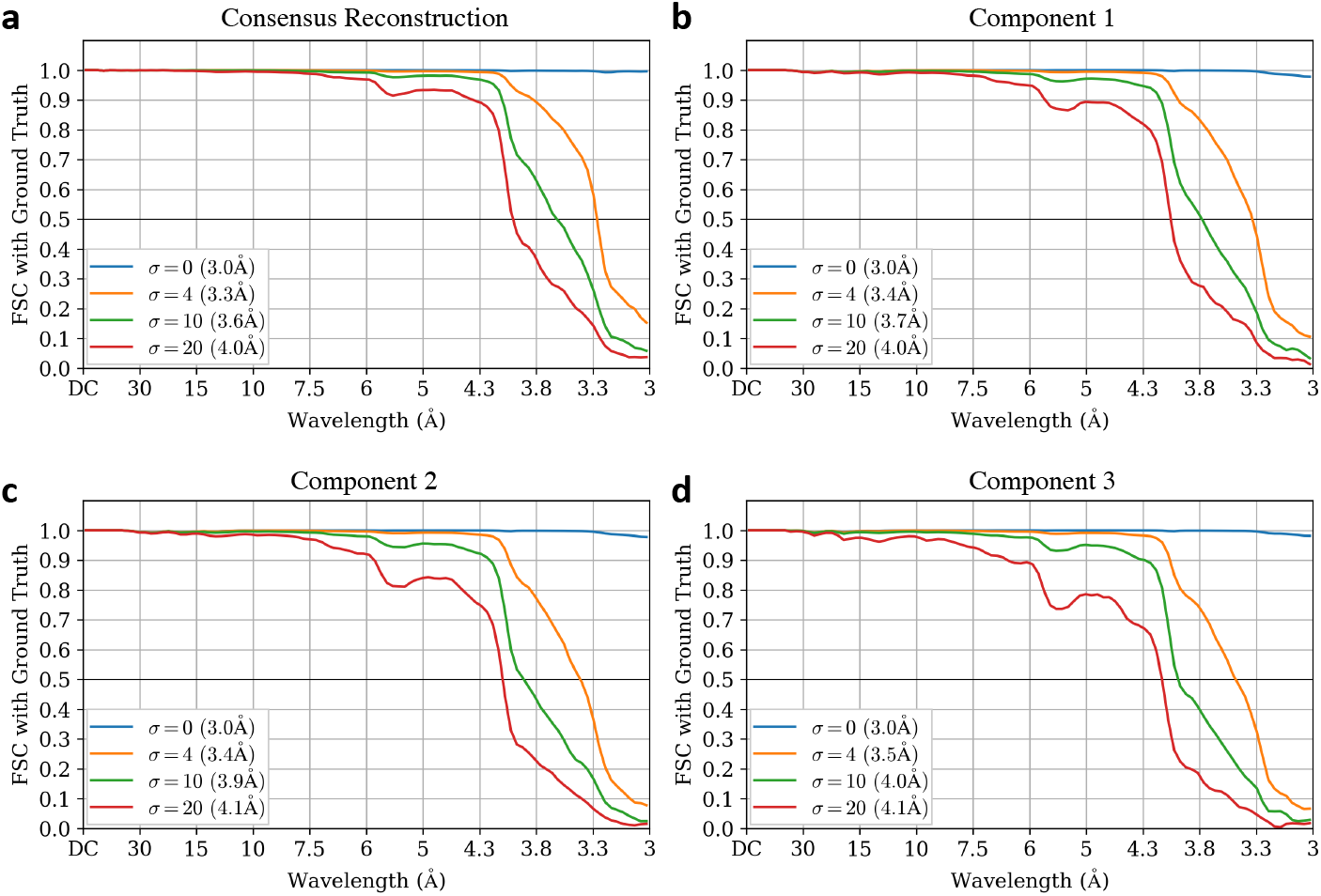
FSC curves of reconstructions with 3DVA to assess resolution of estimated variability components with different noise levels in the synthetic T20S data. **a:** Baseline FSC curves between the ground-truth consensus density map and consensus reconstruction from particle images, at each noise level. **b,c,d:** FSC curves between the ground-truth linear subspace direction and the 3DVA estimated variability component, for components 1, 2, and 3 respectively. In the noiseless case, 3DVA recovers the true subspace directions nearly perfectly, with resolution of variability components decreasing as noise increases.

Next, we compute FSC curves between the 3DVA estimated variability components and the ground truth subspace directions. Figure 4b-d show the FSC curves for components 1, 2, and 3 respectively. Again, we see that in the noiseless case, 3DVA recovers the true subspace directions with nearly perfect correlation at all frequencies. As noise increases, components are resolved with decreasing resolution. Notably, the decrease in resolution of variability components mirrors the decrease in consensus refinement resolution, indicating that the inherent noise in particle images is the main limiting factor for 3DVA estimates, rather than the 3DVA algorithm. In particular, in the noisiest case of *σ* = 20, the consensus reconstruction is limited to an FSC=0.5 value of 4.0*Å* while all three subspace components reach nearly the same resolution, 4.1*Å*.

### 3.2 Cannabinoid GPCR: High resolution flexible motion of small proteins

GPCRs are small membrane proteins responsible for cell signalling and transmission [16]. 3DVA is applied to a dataset of Cannabinoid Receptor 1-G GPCR complex particles [20], containing the CB1 GPCR, G protein, and scFv (single-chain variable fragment scFv16 [20]). Raw microscope images (EMPIAR-10288) are processed in *cryoSPARC* v2 to obtain a consensus refinement of the entire structure (Fig. 5a) from 250,649 particle images. When 3DVA is first run on the entire 3D map, the variability components are dominated by changes in the shape and position of the micelle (Fig. 5b and 5c). To inspect the heterogeneity of the protein itself, a mask is used to exclude solvent and detergent micelle (Fig. 5d). 3DVA is then run using three variability components and a low-pass filter resolution of 3Å.

**Figure 5:**
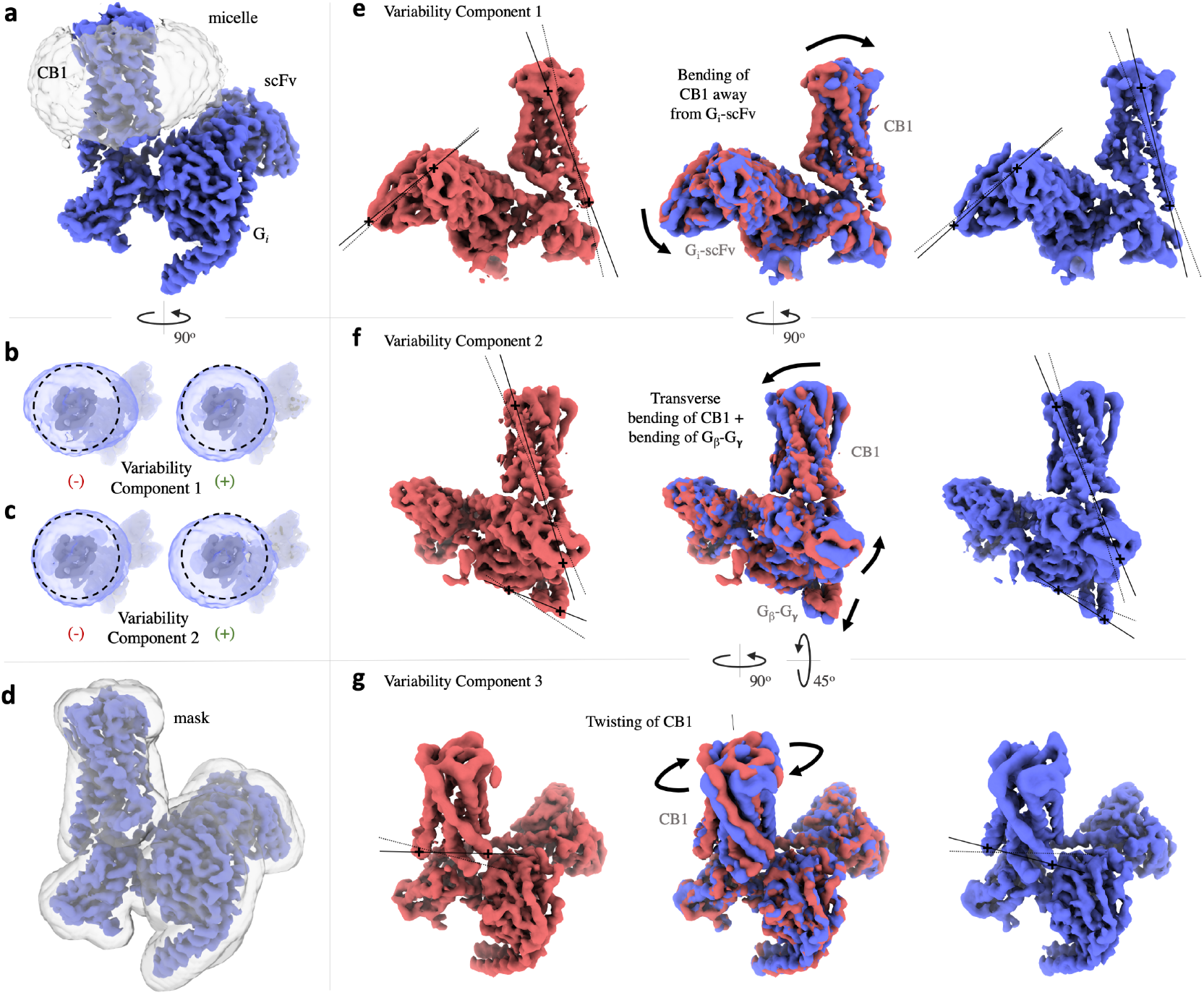
Results of 3DVA with three variability components on 250,649 particles of a Cannabinoid Receptor 1-G GPCR complex [20], demonstrating the capacity for 3DVA to resolve high resolution motion of small proteins and small subregions of proteins. (A video of these results are available in supplementary materials.) **a**: Consensus 3D refinement density of all particle images. The micelle is transparently shaded. **b-c**: Results of 3DVA with two components, with a mask that only excludes solvent. Both components (shown at negative and positive values of latent coordinate) resolve different types of change and motion of the micelle, rather than protein. **d**: Mask that is used for subsequent 3DVA processing that excludes both solvent and micelle. **e**: Component 1 resolves bending of the CB1 transmembrane region away/towards the G-protein. Bending also affects the pose of the scFv subunit. **f**: Component 2 resolves a perpendicular (compared to component 1) bending of the CB1 region, with simultaneous motion of the *G_β_*-*G_γ_* pair of helices. **g**: Component 3 resolves twisting of the CB1 transmembrane domain around an axis perpendicular to the membrane. **e-g**: Markers (+) and guide lines are attached to the same 3D features in negative and positive density maps, as visual reference points.

For the Cannabinoid receptor (Fig. 5e-f), the first component resolves bending of the CB1 transmembrane domain towards and away from the G-protein. The second component resolves a perpendicular bending of the CB1 domain, with simultaneous motion of the *G_β_*-*G_γ_* domain. The third component resolves twisting of the CB1 transmembrane region around a vertical axis, perpendicular to the membrane. This result is notable, as 3DVA is able to resolve the detailed motion of a small subregion of the complex. The entire CB1 protein is only 53 kDa [18] and the CB1 transmembrane domain is even smaller.

3DVA is, to our knowledge, the first method capable of resolving high resolution continuous flexibility for a protein as small as a GPCR complex. An early version of 3DVA in *cryoSPARC* was also used successfully to understand the differences in conformational dynamics between adrenomedullin 1 (AM1) and adrenomedullin 2 (AM2) receptors [22].

### 3.3 80S Ribosome: Solving multiple types of heterogeneity

Applied to 105,247 *Pf* 80S ribosome particles (EMPIAR-10028 [49]), 3DVA with four variability components resolves four types of heterogeneity (Fig. 6). The first component identifies the presence or absence of a portion of the “head” region of the 40S small subunit. The second resolves rotational motion of the entire 40S subunit along an axis that connects the 40S and 60S subunits. The third component resolves lateral motion of the 40S subunit. The fourth resolves transverse rotational motion of the “head” region of the small subunit, perpendicular to the rotation in the second component.

**Figure 6:**
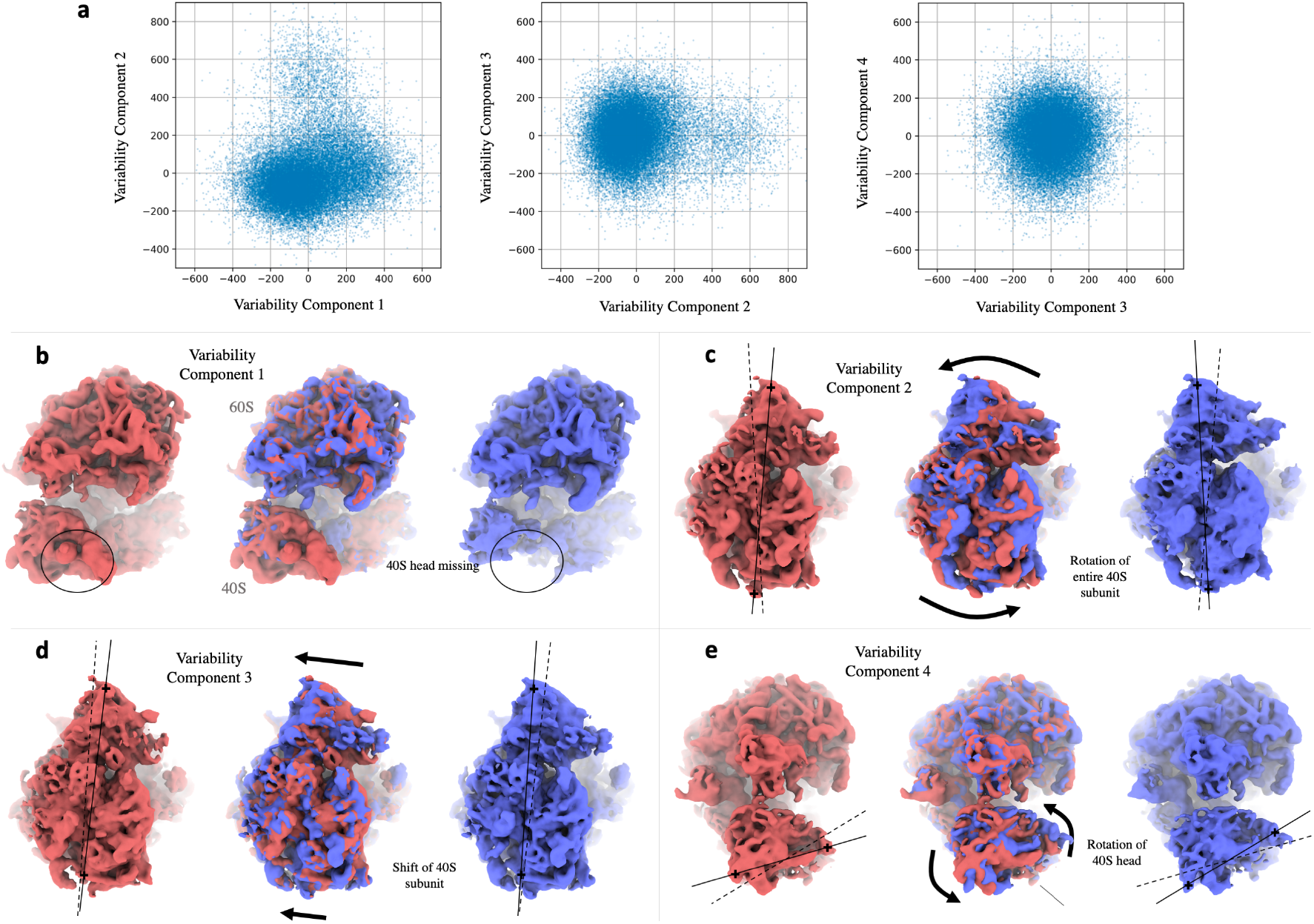
Results of 3DVA with four variability components applied to 105,247 particle images of *Pf* 80S Ribosome [49]. A video of these results is available in supplementary materials. **a**: 2-D scatter plots of 4-D latent coordinates of individual particles are solved by 3DVA. The scatter plots indicate the extent of variability along each dimension. **b-e**: Renderings of 3D density maps generated by 3DVA at negative (red) and positive (blue) positions along each variability component. **b**: (front view) Component 1 identifies the presence and absence of the 40S head region. **c**: (bottom view) Component 2 resolves the rotation of the entire 40S subunit. **d**: (bottom view) Component 3 resolves a lateral shifting of the 40S subunit. **e**: (side view) Component 4 resolves the transverse rotation of the intact 40S head region.

In this case, 3DVA was run with low-pass filter resolution set to 8Å. The four orthogonal types of heterogeneity in the molecule were solved in a single run of 3DVA, using the full resolution particles images, with a 360 pixel box size, taking 71 minutes on a single NVIDIA V100 GPU. As a comparison, on this dataset the recently proposed cryoDRGN deep generative model [53] was reported to resolve conformational changes similar to variability components 1, 2, and 4 [52], but required model training for 150 epochs taking over 60 hours (i.e., 50× more than 3DVA) on the same hardware.

### 3.4 Na_v_1.7: multiple high resolution modes of bending in a sodium ion channel

The Na_v_1.7 channel [51] is a voltage-gated sodium channel found in the human nervous system. Na_v_ channels are fundamental to the generation and conduction of action potentials in neurons. They are mutated in various diseases, and are targeted by toxins and therapeutic drugs (e.g., for pain relief). A dataset of 431,741 particle images of a Na_v_-Fab complex is created by complete processing in *cryoSPARC v2* from raw data (EMPIAR-10261). This particle set is processed with standard 3D classification methods to separate the discrete conformational states of active and inactive channels. The active class contains 300,759 particles. The overall protein-Fab complex is C2 symmetric, and so the particle images are duplicated with their 3D poses rotated 180*°* around the symmetric axis (i.e. symmetry expansion). The resulting 601,518 particles are then input to 3DVA, with six components and a low-pass filter resolution of 3Å.

Of the six components, Figure 7a-c displays three. The first (component 1), resolves bending of two of the transmembrane subunits of the tetrameric protein, along with motion of the bound Fabs. The outer transmembrane helices move left and right while the Fabs move closer and further apart. The second component (component 2), resolves the lateral bending of the 4-helix bundle. The third component (component 6) resolves bending of the two subunits that are not bound to Fabs, in an up-down direction. For this data, refinement resolution of the peripheral transmembrane helices is limited [35]; 3DVA provides insight into the flexibility that causes this limitation.

**Figure 7:**
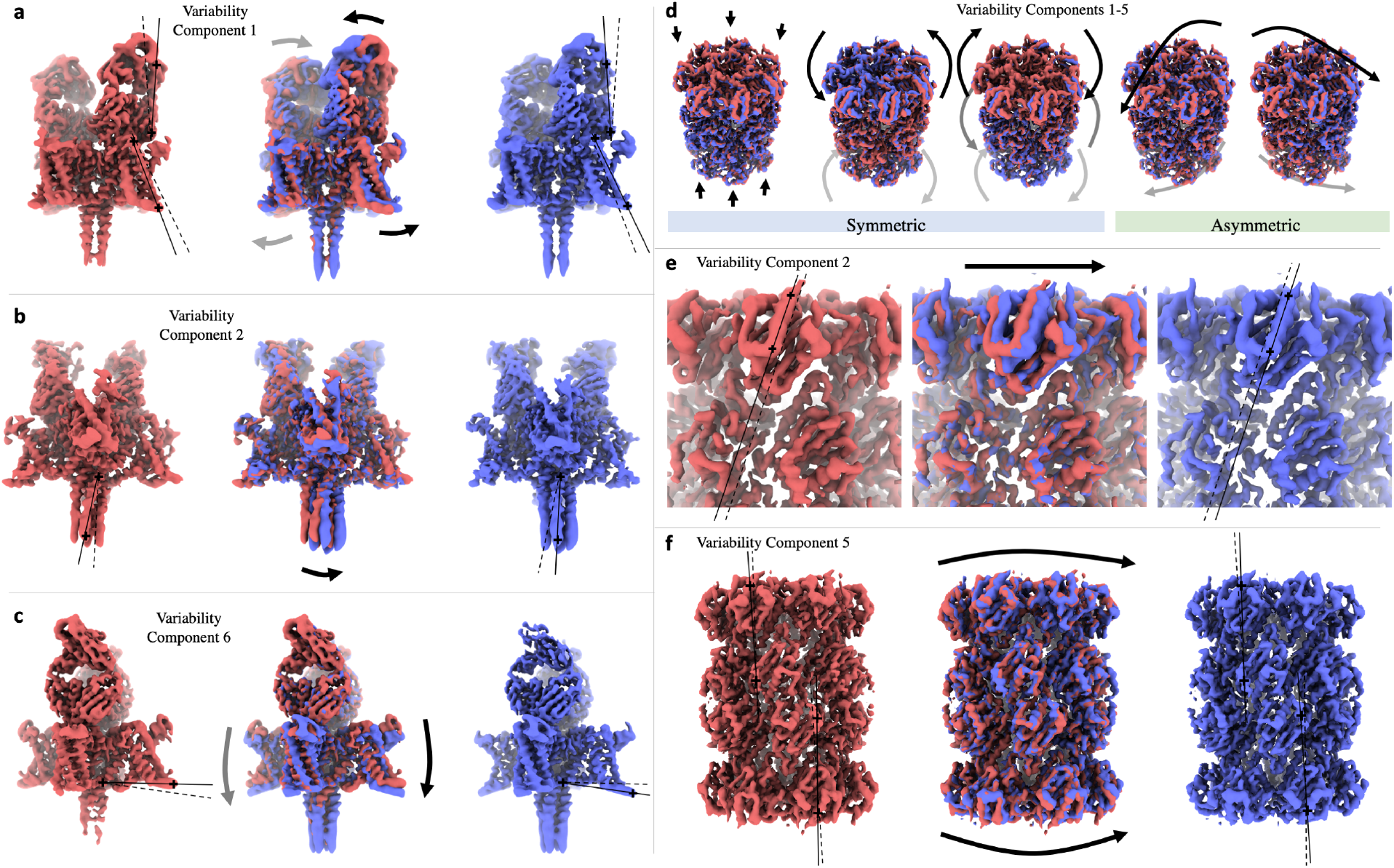
**a-c**: Results of 3DVA with six variability components (three shown) on 300,759 particle images of the Na_v_1.7 channel membrane protein [51]. Particles are subject to symmetry expansion around a C2 symmetry. A video version of these results are available in supplementary materials. **a**: Component 1 resolves simultaneous bending of the two protein subunits that are bound to Fabs, and the motion of the Fabs together/apart. **b**: Component 2 resolves lateral bending of the 4-helix bundle. **c**: Component 6 resolves up/down bending of the two subunits that are not bound to Fabs. **d-f**:Results of 3DVA with five variability components on 84,604 particle images of the T20S Proteasome [4]. Particles are subject to symmetry expansion around a D7 symmetry. A video version of these results are available in supplementary materials. **d**: Overlayed renderings of 3D density maps generated by 3DVA at negative (red) and positive (blue) latent coordinate values along each of five variability components. The first three components resolve symmetric flexible motion of all subunits simultaneously. The next two components resolve asymmetric bending of the entire molecule where the flexing of each subunit is different. **e**: Detail view of variability component 2, showing rotational motion of the top region of the barrel. **f**: Detail view of variability component 5, showing bending of the entire molecule, breaking D7 symmetry.

### 3.5 T20S proteasome: Symmetric and asymmetric flexible motion at high resolution

The T20S proteasome is a D7-symmetric large, stable protein that is commonly used as a test specimen for cryo-EM microscopes [4]. A dataset of 84,605 particle images of T20S is created by complete processing in *cryoSPARC v2* from raw data (EMPIAR-10025). The particle images are duplicated around the 14-fold D7 symmetry (i.e. symmetry expansion), and the resulting 1,184,470 particles are then used for 3DVA with five variability components and a low-pass filter resolution of 5Å.

Despite the generally stable nature of the T20S protein, 3DVA detects five types of continuous bending flexibility in the molecule (Fig. 7d-f). This is an interesting case due to the high symmetry; three of the variability components (Fig. 7d, left) are symmetric, corresponding to symmetric motion of all 14 subunits. The first component resolves a stretching-compression motion of the top and bottom regions of the barrel. The second component (Fig. 7e) shows rotational motion of the top and bottom of the barrel in opposite directions around the symmetry axis. The third component resolves a twisting motion where the middle and ends of the barrel both rotate around the symmetric axis. The last two components are asymmetric (Fig. 7d, right), resolving bending of the entire barrel in two different directions (detail of component 4 in Fig. 7f). Importantly, this shows that 3DVA can resolve several orthogonal modes of detailed high resolution flexible motion of larger molecules. It can also help detect pseudo-symmetries created by asymmetric flexibility of symmetric molecules.

### 3.6 Spliceosome: Large flexible motions of large complexes

On 327,490 particle images of a pre-catalytic spliceosome complex (EMPIAR-10180 [32]), 3DVA was run with two components and a low-pass filter of 8Å, resolving two large motions of multiple parts of the complex (Fig. 8). The first component (Fig. 8b) resolves large rotational motion of both the helicase and SF3b regions towards the front/back of the complex, while the foot region also bends slightly forward and back. The second component (Fig. 8c) resolves side-to-side rotation of SF3b, diagonal rotation of the helicase, and slight side-to-side bending of the foot region.

**Figure 8:**
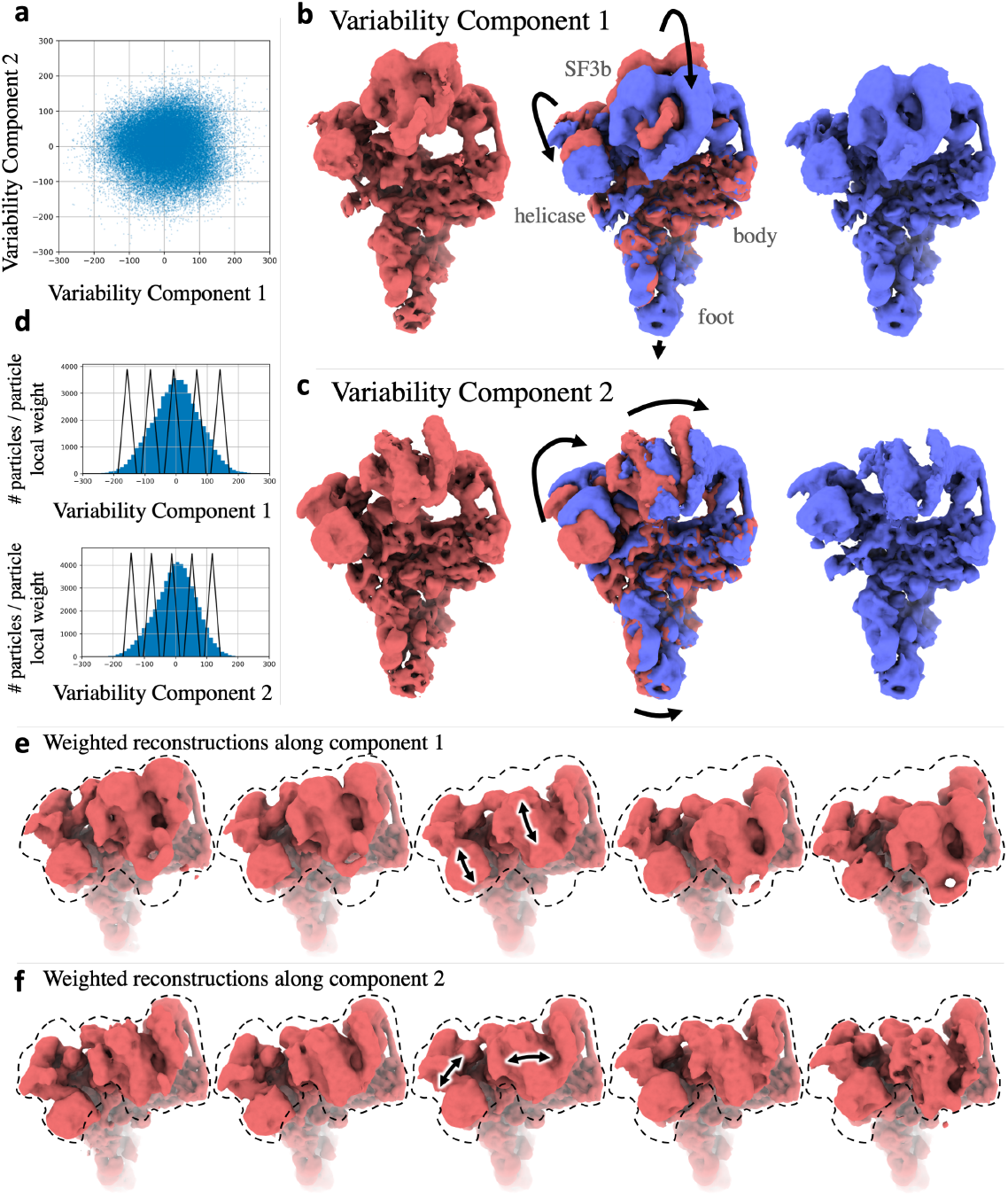
Results of 3DVA with two variabiltiy components on 327,490 particle images of a pre-catalytic spliceosome complex [32]. A video version of these results are available in supplementary materials. **a**: Particle images are spread over two latent coordinates. **b-c**: 3D density maps generated by 3DVA at negative (red) and positive (blue) latent coordinate values. Component 1 resolves forward/back rotation of SF3b and helicase regions, and slight forward/back bending of the food region. Component 2 resolves side-to-side rotation of SF3b and diagonal rotation of helicase. **d**: 1D histograms of both variability components showing local weightings for each of five intermediate positions that are used for weighted reconstructions. **e-f**: Weighted local reconstructions along both variability components, showing five intermediate positions along both types of motion.

The linear subspace model captures both types of large motion, and the latent coordinates of individual particles provide an estimate for the position of the helicase and SF3b regions in each image (Fig. 8a, top). Nevertheless, the linear subspace model underlying 3DVA is limited in its ability to faithfully represent large motions (Fig. 1), so a simple local weighting scheme is used to create intermediate 3D reconstructions along each variability component. Particles are weighted based on their position along each latent coordinate (Fig. 8a, bottom) and 3D reconstructions are created using those weighted particles. This type of local neighborhood weighting is commonly used in manifold embedding applications, and has been used previously in methods for manifold embedding of cryo-EM data [6]. The resulting 3D density maps depict the molecule at different positions along its continuous flexibility (Fig. 8d,e).

The pre-catalytic spliceosome data was processed in 3DVA in 176 minutes. By comparison, on this data, the recently proposed cryoDRGN deep generative model [53] was reported to resolve conformational changes similar to the two variability components [52], but required model training for 30 hours on less than half the number of particles (i.e., 20× slower than 3DVA) on the same hardware.

### 3.7 Bacterial Large Ribosomal Subunit: directly resolving discrete heterogeneity

The bacterial large ribosomal subunit (LSU) is a large macromolecular complex; the cryo-EM dataset contains a mixture of assembly intermediates (EMPIAR-10076 [7]). 3DVA is applied to 131,899 particles, with four variability components and a low-pass filter of 5Å. The 3DVA components lie in a subspace that spans several discrete 3D states. Within that subspace (Fig. 9a) the latent coordinates of particle images are well clustered in multiple dimensions. We find that each cluster corresponds to a different partial assembly state of the complete LSU complex.

**Figure 9:**
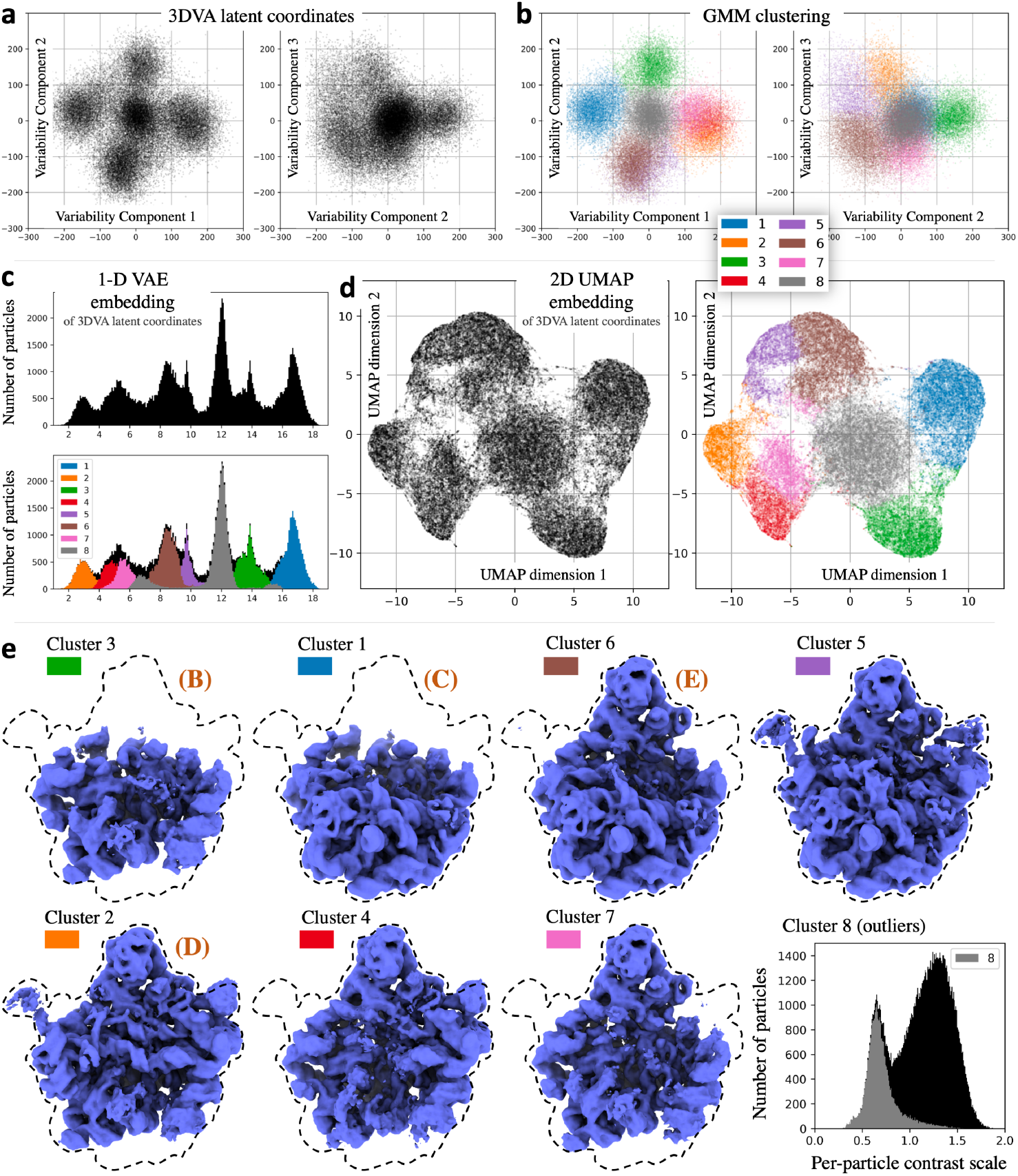
3DVA with four components on 131,899 particle images of a bacterial ribosomal large subunit [7]. 3DVA identifies clusters corresponding to discrete conformational states. **a**: 3DVA particle latent coordinates, as 2D scatter plots of sequential pairs of dimensions. **b**: Latent coordinates colored by 8-way Gaussian Mixture Model (GMM) clustering. **c**: 1D histogram of Variational Autoencoder (VAE) embedding of latent coordinates. The histogram shows seven major peaks. Overlayed histograms (bottom) of each of the GMM clusters from (b) show that peaks correspond to clusters. **d**: 2D UMAP [25] embedding of latent coordinates, showing overall geometry of clusters. Points are colored (right) by GMM clusters from (b). **e**: 3D reconstructions from particles in each GMM cluster. Cluster 3 contains the least assembled complex, and other clusters have more subunits assembled. The 8th cluster, depicted as a histogram showing all particles in black and 8th cluster particles in gray, is composed of outliers for which 3DVA estimates lower per-particle scale factor *α*. This may be because the outliers are broken/denatured particles that do not contain the entire complex.

To analyze and understand the clustering of particles in the 3DVA latent coordinate space, we first fit an 8-component Gaussian Mixture Model (GMM) to the latent coordinates (Fig. 9b). This clusters particles into 8 major groupings. The particles from each cluster are then used to perform a 3D reconstruction; Fig. 9e shows the resulting 3D densities. Cluster 3 contains the least density, and each other cluster adds on a part to the assembly. Cluster 5 displays the entire LSU complex. Clusters 1, 2, 3 and 6 correspond to the four large 3D class subpopulations reported in the original study (C, D, B, E respectively) [7]. Cluster 8 appears to contain mainly outlier particles (contaminants and junk particles). The presence of outliers was also noted in the original study [7]. The identity of this cluster as outliers can be seen in the histogram of estimated per-particle contrast scales *α* in Fig. 9e, where particles in cluster 8 have a low estimated contrast (indicating a low level of agreement between the particles and the consensus 3D density) compared to all other clusters.

To further investigate and demonstrate the power of 3DVA components to capture discrete heterogeneity, two non-linear embedding methods are applied to the 4-D per-particle latent coordinates. First, a Variational Autoencoder (VAE) [19] is trained to reduce the 4-D latent coordinates to a 1-D embedding where individual clusters can be directly visually identified. The VAE is constructed using *PyTorch* [30], with a single hidden layer of size 1024 in both encoder and decoder architectures. Training took 2 minutes for 40 epochs. The resulting 1-D latent encoding is shown in Fig. 9c as a histogram of particle positions. The peaks correspond to the multiple discrete states and the outlier cluster detected by GMM fitting (with the same colors as Fig. 9b). This result can be directly compared with the 1-D Spatial-VAE embedding carried out in processing of this same dataset using the cryoDRGN generative model [52], where a similar histogram is generated that resolves four major sub-states, plus an outlier cluster.

A second non-linear embedding technique, UMAP [25], is applied to the 4-D latent coordinates (Fig. 9d), displaying the geometry of clusters in the 4-D space, e.g., that clusters 2, 4, and 7 are sub-clusters of a larger cluster. This can also be compared with results in [52] to illustrate the ability of 3DVA to resolve nuanced discrete conformational heterogeneity, despite the apparent simplicity of the linear subspace model.

## 4 Discussion

The 3DVA algorithm enables one to fit high-resolution linear subspace models to single particle cryo-EM data. The output variability components and associated latent coordinates reveal biological insights from the data, including detailed flexible motion at the level of individual *α*-helices, even for small membrane proteins. It can can also be used to identify discrete conformational states of proteins from heterogeneous samples.

### Related methods for manifold fitting

Linear subspace models, such as PCA, Multivariate Statistical Analysis (MSA), and eigenanalysis have been explored for single particle cryo-EM. Early methods used bootstrapping 3D reconstructions from random subsets of images to coarsely simulate sampling from the conformational distribution of a target molecule [31]. Others attempt to construct a low-resolution complete covariance matrix of the 3D density map distribution underlying a sample, extracting a linear subspace using eigenanalysis [1].

Tagare et al. [46] proposed a similar subspace model, using maximum likelihood estimation with a form of Expectation-Maximization. However, their M-step is solved via an approximate iterative conjugate gradient, and each *V_k_* is solved sequentially, requiring *K* complete rounds of optimization, each passing through the data repeatedly until convergence for each *V_k_*. Furthermore, the CTF is treated approximately using Wiener filtering, image noise is assumed to be white, and particles are binned into a finite number of 3D pose bins. Thus, they were only able to compute linear subspace models for coarse low-resolution motion and variations; results on experimental data are demonstrated only at 15Å resolution, and only for *K* = 2 variability components. 3DVA computes the M-step exactly, without approximate methods, and all *K* variability components are solved simultaneously at full resolution, and the per-particle scale factors *α_i_* and a colored noise model are optimized in parallel.

Non-linear manifold embedding techniques are also in development. Diffusion maps have been used in 2D, with multiple 2D manifold embeddings combined to resolve a 3D manifold of continuous conformational change [6, 10]. These methods have been demonstrated on experimental data of large proteins, and recent work aims to improve the difficult process of combining 2D manifolds to form a 3D model [24]. Non-linear methods employing the graph Laplacian of similar 2D particle images have also been developed [27] and applied to synthetic data. Most recently, non-linear, deep generative models, have been proposed and shown to resolve coarse-scale conformational changes from experimental cryo-EM data of large complexes [52, 53]. Methods that directly model protein motion have also been proposed, but only demonstrated on simplified synthetic data [12, 21].

### Relationship to other SPA heterogeneity techniques

The predominant method currently used to resolve heterogeneity in cryo-EM data is discrete 3D classification [13, 40, 42, 43]. By comparison, a linear subspace model does not constrain particles to exist in a (small) finite number of conformational states. Rather, the subspace components span a continuous space of possible states. It is well-known in the cryo-EM field that clustering methods are not effective for resolving continuous flexibility.

Local refinement [13, 42] and multi-body refinement [28] methods allow high resolution refinement of some regions of flexible molecules, by assuming the molecule is composed of a small number of rigid parts. For successful 2D-3D alignment, such methods require that each region has sufficient mass for accurate local alignment, independent of the rest of the structure. 3DVA does not make this assumption, as it allows generic continuous flexibility within the subspace.

Techniques like normal-modes analysis [5] make assumptions about the energy landscape and dynamics of a protein molecule to predict possible flexibility. Methods have been proposed to exploit these fixed prior models to recover information from cryo-EM data of flexible molecules [44]. In contrast, 3DVA does not presuppose knowledge of the energy landscape, bending, or flexing of the molecule. Rather, it learns this from the image data.

Finally, linear subspace models have also been useful in cryo-electron tomography (cryo-ET), specifically in sub-tomogram averaging where multiple 3D reconstructions of individual molecules, are obtained. By averaging such reconstructions one can improve resolution. In this case, PCA can be directly applied [15, 17] since tomography provides complete 3D observations for each individual particle.

### Properties of 3DVA

3DVA has several properties that make it particularly attractive and useful. First, 3DVA is not sensitive to initialization bias. Generally, iterative algorithms for non-convex optimization problems can give incorrect results if initialized poorly. It has been proven, however, that for PPCA models (like 3DVA) any stable local optima is also a global optimum of the objective function [47]. This means that no matter how *V*_1:*K*_ are initialized at the start of optimizing 3DVA, the results will be equivalent. This is in contrast to existing methods for discrete 3D classification and other non-linear methods, where initialization is critical and multiple runs often yield different results.

It also notable that 3DVA, using Expectation-Maximization, is a Krylov subspace method [39]. Krylov subspace methods are a general class of iterative optimization algorithms for computing the eigenvectors of matrices, used in popular linear algebra techniques like SVD [11]. Krylov subspace methods like 3DVA have the beneficial property of solving eigenvectors in order of importance. Running 3DVA with *K* = 3 will yield the top three variability components that explain the most variability in the data, and running with *K* = 6 will yield the same three plus the next three important components.

With these properties and the stability of Expectation-Maximization, 3DVA does not require tuning or parameter changes between datasets, beyond setting the resolution limits and the number of components, *K*. Since 3DVA depends on the quality of the mean density, *V*_0_ and the particle alignments, methods that produce high quality consensus structures and alignments in the presence of flexibility and heterogeneity, e.g., non-uniform refinement [35], can improve the results of 3DVA.

### Extensions and next steps

The 3DVA algorithm uses a linear subspace model of 3D structures that comprise the conformational landscape of a protein molecule. The true manifold can have non-linear and complex geometry, for which non-linear manifold models are likely required. For example, extensions of 3DVA using similar algorithmic techniques may be possible that allow for some forms of non-linearity. The development of more accurate and powerful generic models of conformational manifolds is only the first step in truly solving flexible protein structures. In addition to modelling the manifold, techniques must be developed that aggregate structural information across continuous conformational states along a manifold, to solve higher resolution 3D density maps of flexible molecules.

## 5 Methods

3DVA is formulated as a form of probabilistic PCA, and solved with the Expectation-Maximization algorithm. Accordingly, before explaining 3DVA in detail, we first review the probabilistic PCA model and the Expectation-Maximization algorithm for parameter estimation. We then provide the formulation of 3DVA, along with the numerical details of the E- and M-steps of the iterative method.

### Probabilistic PCA and Expectation-Maximization

The PCA model, in abstract terms, specifies that an observed data point (e.g., vector **x**) is equal to some mean vector *μ* plus a linear mapping *B* from a latent coordinate vector **z**:

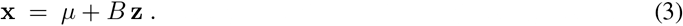

To solve for the unknowns, i.e., *B*, *μ* and **z**, one sets *μ* to be mean of the observed data, and the columns *B* are the leading eigenvectors of the covariance matrix of the observed data. The latent coordinate vector **z**_*j*_ corresponding to data point **x**_*j*_ is then equal to *B^T^* **x**_*j*_.

Probabilistic PCA (PPCA) is a probabilistic extension of PCA [39, 47]. The PPCA model assumes the latent coordinate vectors are Gaussianly distributed, with zero mean and a unit diagonal covariance matrix, denoted by 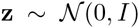, where *I* is the identity matrix. PPCA also assumes additive Gaussian noise in the generation of data, **x** = *μ* + *B***z** + *η*, where *η* is also assumed to be Gaussian, zero-mean and isotropic, with variance *ϵI*; i.e., 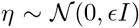. In a seminal paper, Roweis [39] showed how to solve for the PPCA unknowns (i.e., *B*, *μ*, *ϵ*, and the latent coordinates for the *J* data points, denoted **z**_1:*J*_) using the iterative Expectation-Maximization algorithm [8, 29]. Each iteration comprises an E-step, which finds the mean and covariance of the Gaussian posterior distribution over latent coordinates, and an M-step that finds maximum likelihood estimates for the remaining parameters [39].

Roweis’ result is important for 3DVA in several ways. First, the maximum likelihood estimator in the M-step permits partial observations, as in cryo-EM where our observations are 2D images, even though the unknowns of interest are 3D density maps. Second, Roweis shows that PCA is a special case of PPCA (occurring when *ϵ* → 0), and that both PPCA and PCA share the same principal subspace. This is useful because the E-step can then compute maximum likelihood estimates for the latent coordinates without the extra cost of estimating the covariance matrices for the latent coordinates. And third, the computational cost of Expectation-Maximization is much lower than conventional PCA, which depends on the square of the dimension of the data in forming the covariance matrix and finding its eigenvectors. In the context of 3DVA, for a 3D box with linear dimension *N*, the data dimension is *N* ^3^, and the cost of PCA scales with *N* ^6^, while the cost of 3DVA with *K* variability components scales with *N* ^3^*K*. Since *N* ^3^ ≫ *K* this difference can be critical.

### 3D Variability Analysis

The generative model for 3DVA is given in Eq. (2). For *J* particles images, we assume we are given a consensus reconstruction *V*_0_ (in Fourier space), along with the per-particle poses and CTFs parameters. The goal is to estimate *K* variability components, *V*_1:*K*_, and the scalar multipliers and latent coordinates for each of *J* particles images, denoted *α*_1:*J*_ and **z**_1:*J*_.

Under the 3DVA model the latent coordinates are assumed to be mean-zero Gaussian with an isotropic covariance matrix; i.e., 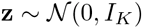, where *I_K_* is the *K × K* identity matrix. The noise is assumed to be zero-mean Gaussian and independent of **z**, and with a diagonal covariance matrix in which the variance is constant for Fourier coefficients in annular rings. Such ‘colored’ noise models are useful with cryo-EM data as they capture the dependence of signal-to-noise ratios on wavelength [34].

Fortunately we can whiten the noise and thereby simplify the formulation. To this end, given the inverse covariance matrix of the noise, denoted by Λ, one can multiply the image *X* and the CTF by Λ^1/2^, yielding 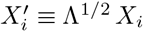, 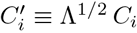, and a generative model with isotropic zero-mean noise *η′*:

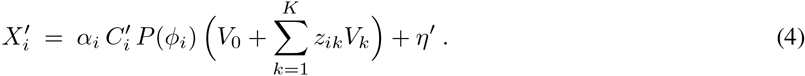

As such, the PPCA model as described in [39] can be adapted for cryo-EM.

More specifically, the 3DVA algorithm is derived from the PPCA model in [39] under the assumption that the noise variance *ϵ* tends to zero, in which case PPCA reduces to PCA. While it may seem counter-intuitive to assume that noise tends to zero for cryo-EM data, it has computational benefits. As shown in [39], both PPCA and PCA share the same linear subspace. As a consequence, the variability components recovered by 3DVA are unchanged by this assumption. Nevertheless, in the limiting PCA case, the E-step only requires that we compute the posterior mean, avoiding the need to estimate the posterior covariance. Further, the M-step reduces to solving a simpler maximum likelihood estimation problem.

It follows that both the E- and M-steps used to fit the subspace model in 3DVA entail the minimization of a single energy function equal to the sum of squared residual errors between the observed images, *X_i_*, and model predictions. Given *J* particle images, the resulting energy function, expressed in the Fourier domain, is given by

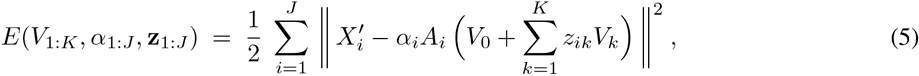

where 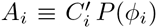. The E-step and M-step can be viewed as a form of iterative coordinate descent. The E-step minimizes the energy in Eq. (5) to find the posterior mean latent coordinates for each particle image. The M-step then decreases the energy with respect to the *K* variability components *V*_1:*K*_ and the per-particle scale factors *α*_1:*J*_

The main challenges with the optimization stem from the large number of unknowns. The E-step computes the posterior means for the latent coordinate, separately for each particle image, given the variability components *V*_1:*K*_ and the scale factors *α_i_*. To this end, let *W_ik_* = *A_i_V_k_* be *k*th variability component after being projected onto 2D, given the pose of the *i*th particle image, and after application of the CTF. Then, the energy for the *i*th image is

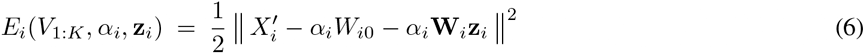

where **W**_*i*_ ≡ [*W_i_*_1_*, …, W_iK_*] and **z**_*i*_ ≡ (*z_i_*_1_, …, *z_iK_*)^*T*^. The columns of **W**_*i*_ are the projections of the *K* variability components according to the CTF and pose of the *i*th particle. Each column has *N* ^2^ elements, the number of Fourier coefficients in *X_i_*. So **W** is a *N*^2^ × *K* matrix, and _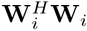_ is a *K × K* symmetric positive-definite matrix, where **W**^*H*^ denotes the conjugate transpose of **W**. The mean latent coordinates is given in closed form by

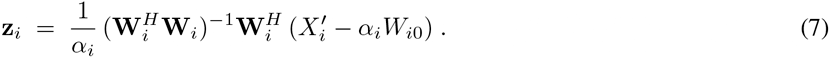

The first phase of the M-step minimizes the energy associated with each particle image in Eq. 6 to solve for the updated scale factors (given the most recent latent coordinates). This is given by

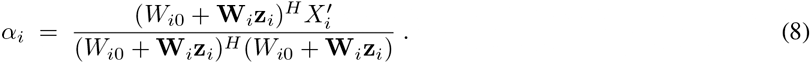

The second computation in the M-step involves the estimation of the variability component updates. To this end it is useful to rewrite the energy, now summed over all images, in a somewhat simpler form:

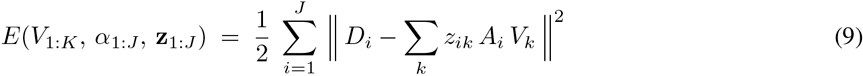

where 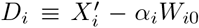 and *z_ik_* is the *k*th latent coordinate for the *i*th particle image. To solve for variability components, we require the gradient of *E* with respect to each *V_j_*:

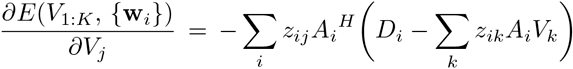

Setting the gradient to zero and rearranging terms yields a linear system of equations,

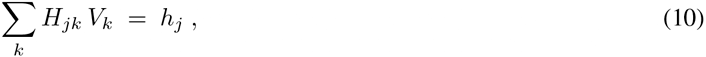

where 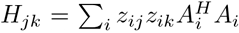 and 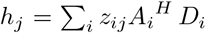. Importantly, *H_jk_* can be represented as a diagonal matrix; to see this, note that 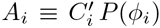, and the CTF operator ^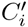^is diagonal. Further, depending on the form of the interpolation used, the projection matrix inner product _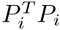_ is either diagonal or approximately diagonal. As is standard in single-particle EM reconstruction algorithms, we approximate this product as a diagonal matrix.

Differentiating *E* with respect to all variability components, we obtain a block matrix system of equations:

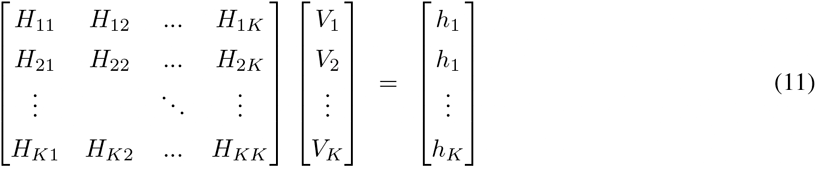

Exploiting the diagonal structure of individual blocks, *H_jk_*, we rearrange the rows and column of Eq. (11) to form a block diagonal system having *N*^3^ decoupled, *K × K* linear systems, one for each wavenumber. Each *K × K* systems is used to estimate the Fourier coefficients at a specific wavenumber for each of the *K* variability components.

To rearrange the left hand side of Eq. (11), the first *K* columns will contain the first column from each of the *K* blocks in the large block *H* matrix in Eq. (11). The next *K* columns comprise the second column of each of the *K* blocks and so on. The rows of the large variability component vector are rearranged correspondingly. This places the Fourier coefficients associated with the first wavenumber together, followed by those associated with the second wavenumber next, and so on. Next we rearrange the rows of the matrix and the corresponding right hand side elements. The first *K* rows comprise the first row from each of the *K* blocks of the *H* matrix on the left hand side of Eq. (11). The next *K* rows comprise the second row of each block, and so on. We also rearrange the corresponding elements on the right hand side. This rearrangement yields a block diagonal system, decoupling the estimation of different wavenumbers.

Finally should we want to impose orthogonality constraints, we could use Lagrange multipliers or constrained optimization, but with Expectation-Maximization it is sufficient to orthogonalize the variability components at the end of each iteration. It is done with the Gram-Schmidt algorithm [11], which is efficient as it only requires inner products between variability components, and scaling each variability component to have unit length, i.e., 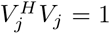. While not technically required by the iterative solution of a linear subspace model, orthogonality improves computational stability by removing degrees of freedom to which the solution is invariant.

In addition to orthogonality, in practice it is often helpful to restrict analysis to a specific region of 3D density using a mask, in real-space. Masking is implemented in each iteration at the end of the M-step, by transforming each variability component into real-space, multiplying by the soft input mask, and transforming back to the Fourier domain. This ensures that any variability outside the masked region is ignored in the subsequent E-step.

Iterations continue until convergence, or for a fixed number of iterations. The experiments here used 20 iterations. Once completed, we form the *K×K* covariance matrix of latent coordinates, 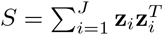. The eigenvectors of *S* specify a rotation of the variability components and the latent coordinates that align the variability components with the principal directions of the data and ensures the latent coordinates are independent under the Gaussian model, as with PPCA. The per-particle scale factors *α_i_* are unchanged under this rotation.

## Supporting information

Movie S1 80S Ribosome

Movie S2 CB1 GPCR

Movie S3 Nav17 Channel

Movie S4 T20S Proteasome

Movie S5 Spliceosome

## Code Availability

The cryoSPARC software package is available for non-profit academic use at www.cryosparc.com.

## Acknowledgements

We thank the entire team at Structura Biotechnology Inc. that designs, develops, and maintains the *cryoSPARC* software system, in which this project was implemented and tested. Resources used in this research were provided, in part, by the Province of Ontario, the Government of Canada through NSERC and CIFAR, and companies sponsoring the Vector Institute.

## Author Contributions

A.P. developed the method, created the implementation and performed experiments. A.P. and D.J.F. created the formulation and wrote the paper.

